# A nuclear import pathway exploited by pathogenic noncoding RNAs

**DOI:** 10.1101/2021.12.02.470923

**Authors:** Junfei Ma, Shachinthaka D. Dissanayaka Mudiyanselage, Woong June Park, Mo Wang, Ryuta Takeda, Bin Liu, Ying Wang

## Abstract

The prevailing view regarding intracellular RNA trafficking in eukaryotic cells describes that RNAs transcribed in the nucleus either stay in the nucleus or cross the nuclear envelope entering the cytoplasm for function. Interestingly, emerging evidence illustrates numerous functional RNAs trafficking in the reverse direction from the cytoplasm to the nucleus. However, the mechanism underlying the RNA nuclear import has not been well elucidated. Viroids are single-stranded circular noncoding RNAs that infect plants. Using nuclear-replicating viroids as a model, we showed that cellular Importin alpha-4 is likely involved in viroid RNA nuclear import, empirically supporting the involvement of Importin-based cellular pathway in RNA nuclear import. We also confirmed the involvement of a cellular protein (Virp1) that binds both Importin alpha-4 and viroids. Furthermore, a conserved C-loop in nuclear-replicating viroids is critical for Virp1 binding. Disrupting C-loop impairs Virp1 binding, viroid nuclear accumulation and infectivity. Further, C-loop exists in a subviral satellite noncoding RNA that relies on Virp1 for nuclear import. These results have significant implications for understanding the infection process of subviral agents. In addition, our data outline a cellular pathway responsible for the nuclear import of RNAs and uncover a 3-dimensional RNA motif-based regulation over RNA nuclear import.

## Introduction

Since most cellular RNAs are produced through transcription in the nucleus of eukaryotic cells, the prevailing view is that those cellular RNAs either stay in the nucleus or move to the cytoplasm for function. Interestingly, emerging evidence showed that cellular RNAs (i.e., small RNAs, tRNAs, and rRNAs), as well as viral RNAs, can traffic in a reverse direction from the cytoplasm to the nucleus. For instance, plant 24-nt hc-siRNAs are exported to the cytoplasm for Argonaut 4 loading before being redirected into the nucleus of the same cell or even neighboring cells for RNA-directed DNA methylation (Ye et al., 2012; Long et al., 2021). In *Xenopus* oocytes, 5S rRNA relies on ribosomal protein L5 for nuclear import (Rudt and Pieler, 1996). In another example, satellite RNA of Q-strain cucumber mosaic virus (Q-satRNA) relies on a bromodomain-containing cellular protein (Virp1) for entering the nucleus (Chaturvedi et al., 2014). By and large, in contrast to the well-studied RNA nuclear export processes, the RNA nuclear import machinery and mechanism remain obscure, particularly regarding the molecular basis underlying the specific selection of RNAs for nuclear import.

To cross the double-membrane nuclear envelope, biomolecules need to traffic through the highly organized nuclear pore complexes (NPCs) in eukaryotic cells (Merkle, 2011). NPCs are conserved in eukaryotic organisms with some variations (Xu and Meier, 2008; Meier and Brkljacic, 2009). Except for some free-diffusing small molecules below 40 to 60 kD, most biomolecules rely on nuclear transport receptors (NTRs) to traffic through NPCs (Frey et al., 2006; Stewart et al., 2007; Frey and Gorlich, 2009; Merkle, 2011). Importin alpha subunits (IMPas) constitute a group of adapter proteins linking specific cargos to NTRs for crossing NPCs (Merkle, 2011). In *Arabidopsis thaliana*, nine IMPas play distinct yet partially redundant roles (Merkle, 2011; Chen et al., 2020). Whether any IMPa is involved in RNA nuclear import remains to be determined.

Viroids are single-stranded circular noncoding RNAs that infect plants (Wang, 2021). Due to their noncoding nature, viroids must utilize RNA structures to exploit cellular factors in order to accomplish their infection cycles. RNA secondary structures are primarily composed of helices and loops. RNA loops often contain highly arrayed non-Watson-Crick (non-WC) base-pairings and other base-specific interactions, including base stacking and base-backbone interactions (Wang et al., 2018). Each RNA base can use its three edges (i.e., WC, Hoogsteen, and sugar edge) to form non-WC base-pairing geometries within an structural motif (Wang et al., 2018). Those non-WC base-pairings have been well documented by a large amount of atomic-resolution crystallography and NMR spectroscopy data (deposited in Protein Data Bank; https://www.rcsb.org). Several homology-based programs have been developed to search for possible base-pairing geometry of a motif of interest (Sarver et al., 2008; Zirbel et al., 2015). The RNA Basepair Catalog summarizes all possible non-WC base-pairings and their similarities from the deposited structural data (Stombaugh et al., 2009), providing a valuable resource for analyzing non-WC base-pairings and predicting functional substitutions (Wang et al., 2018). Such an approach, in combination with functional mutagenesis, has been successfully applied to analyze the structure-function relationships of multiple viroid motifs (Zhong et al., 2006; Zhong et al., 2007; Takeda et al., 2011; Takeda et al., 2018; Wu et al., 2019).

Viroid RNA secondary structures have been well annotated via various chemical mapping assays (Gast et al., 1996; Xu et al., 2012; Giguere et al., 2014; Giguere and Perreault, 2017), providing a solid foundation to annotate base interaction geometries within loop motifs. Recent genome-wide analysis on potato spindle tuber viroid (PSTVd) RNA motifs has identified 11 out of 27 loop motifs responsible for systemic infection (Zhong et al., 2008). Some of those loop motifs regulate RNA trafficking across certain cellular boundaries, and their 3-dimensional (3D) structures have been successfully annotated using a combination of program prediction and functional mutagenesis (Zhong et al., 2007; Takeda et al., 2011; Takeda et al., 2018; Wu et al., 2019). However, whether any RNA motif regulates viroid subcellular localization and organelle targeting remains unknown. Viroids of the family *Pospiviroidae* all replicate in the nucleus, and their nuclear import process is highly regulated (Woo et al., 1999; Seo et al., 2020). Hence, their noncoding RNA genomes likely contain the necessary information in certain forms (e.g., an RNA 3D motif) to guide nuclear import. The cellular factor(s) for viroid nuclear import remains elusive as well. One viroid binding protein, Virp1, has been implied to accelerate the import of citrus exocortis viroid (CEVd) to nuclei of onion cell strips (Seo et al., 2021). Nevertheless, whether and how Virp1 regulates viroid nuclear import await to be clarified.

To gain a better understanding of RNA nuclear import, we test the viroid interaction with all nine IMPas cloned from *Arabidopsis* via RNA-immunoprecipitation and identified IMPa-4 as the specific factor interacting with our model viroid, potato spindle tuber viroid (PSTVd). Down-regulation of the IMPa-4 ortholog in tomato led to a drastic reduction of PSTVd infection. We also showed that IMPa-4 interacts with Virp1 in vivo via co-immunoprecipitation. Moreover, we found that Virp1 recognizes a specific RNA 3D motif, C-loop, in PSTVd and hop stunt viroid (HSVd). Mutational analyses showed that viroid C-loop is critical for Virp1 binding, viroid nuclear accumulation and infectivity. Notably, C-loop can be found in nearly all the nuclear-replicating viroids and also in the Q-satRNA that relies on Virp1 for nuclear import. This work significantly advances our understanding of subviral RNAs. In addition, our data unravel a cellular pathway for RNA nuclear import and the molecular basis of a nuclear import signal in RNAs, which have profound implications in understanding the intracellular trafficking of viral RNAs, and potentially cellular RNAs as well.

## Results

### IMPa-4 is responsible for PSTVd nuclear import

To uncover the specific IMPa protein(s) for PSTVd nuclear import, we employed the RNA-immunoprecipitation assay to determine the PSTVd-binding ability of nine IMPa proteins cloned from *A. thaliana*. As shown in Figure 1A, only IMPa-4 specifically and consistently interacted with PSTVd, as revealed by the presence of PSTVd in the immunoprecipitated fractions via RT-PCR. We chose *Histone H2A* mRNA (*Niben101Scf01866g00004*.*1*) as a negative control for RT-PCR because mRNAs cannot traffic back to the nucleus. Moreover, the H2A ortholog in tomato did not change expression level in PSTVd- or virus-induced gene silencing vector (tobacco rattle virus; TRV)-infected plants in our previous studies (Zheng et al., 2017a; Zheng et al., 2017c). As shown in Figure 1A, IMPa-4 did not bind with the *Histone H2A* mRNA, further supporting the specificity in IMPa-4/PSTVd binding. Homology-based analysis showed that IMPa-4 is also present in tomato (Supplemental Table 1), a host of PSTVd. None of the tomato IMPas, including IMPa-4, have any significant change in expression in PSTVd-infected leaves based on our previously published RNA-Seq data (Supplemental Table 1).

**Figure 1.**
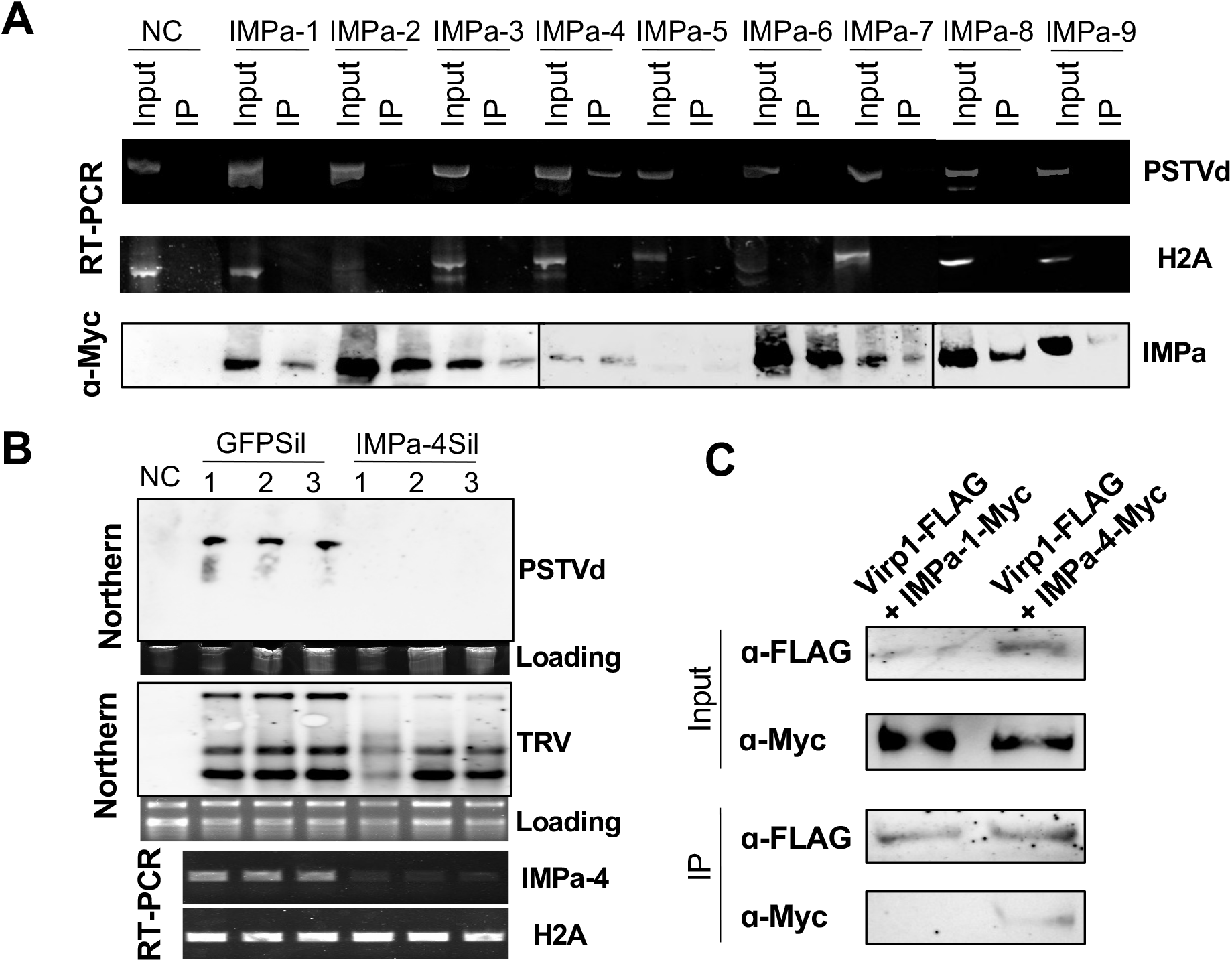
IMPa-4 and Virp1 in the same pathway for PSTVd infection. (A) RNA-immunoprecipitation. TAP-tagged IMPa genes were used to pull down RNAs, which were detected by RT-PCR. Histone H2A serves as a negative control. NC, negative control. IP, immunoprecipitated fraction. (B) RNA gel blots showing PSTVd and TRV accumulation in infiltrated leaves. RT-PCR showing the specific downregulation of IMPa-4 by the TRV-IMPa-4Sil construct. C) Co-immunoprecipitation. FLAG-tagged Virp1 serves a bait to pull down IMPa proteins. IP, immunoprecipitated fraction. H2A, histone H2A. NC, negative control.

No viroid is known to infect *A. thaliana* systemically. Therefore, we employed the virus-induced gene silencing assay to specifically down-regulate the IMPa-4 ortholog in tomato and tested PSTVd infection therein to corroborate the role of IMPa-4 in PSTVd infection, The TRV-GFPsil served as a control that did not affect PSTVd infectivity (Figure 1B). We cloned an IMPa-4-specific fragment based on the BLAST result and constructed TRV-IMPa-4sil. As expected, the TRV-IMPa-4sil construct strongly suppressed PSTVd accumulation in systemic leaves (Figure 1B). In addition, IMPa-4 expression was suppressed in TRV-IMPa-4sil but not TRV-GFPsil-treated leaves (Figure 1B). These data confirmed that IMPa-4 indeed facilitates viroid nuclear imports in plants.

### IMPa-4 interacts with Virp1

Virp1 was discovered through screening a cDNA library from PSTVd-infected tomato for RNA ligand binding (Martinez de Alba et al., 2003) and shown to affect viroid trafficking (Maniataki et al., 2003) and replication (Kalantidis et al., 2007). Down-regulation of Virp1 is known to attenuate viroid replication in cells (Kalantidis et al., 2007). Recent progress showed that Virp1 is responsible for the nuclear import of Q-satRNA (Chaturvedi et al., 2014). However, whether Virp1 is responsible for viroid nuclear import remains elusive. If Virp1 is critical for viroid nuclear import, it likely functions in the same pathway as IMPa-4. To test this possibility, we employed the co-immunoprecipitation assay to test the interaction between these two proteins. We co-expressed a FLAG-tagged Virp1 construct with TAP-tagged IMPa-4 or IMPa-1. As shown in Figure 1C, Virp1 only interacted with IMPa-4 but not IMPa-1. Therefore, Virp1 and IMPa-4 likely form a complex and function in the same pathway.

### A 3-dimensional RNA motif mediates Virp1 binding with PSTVd

Previous analysis suggested that Virp1 binds to two possible RY motifs in PSTVd (Gozmanova et al., 2003), but the structural basis of the RY motif remains elusive. Furthermore, despite that a similar RY motif has been found in another nuclear-replicating viroid HSVd, the overall structures of the RY motif-containing regions between PSTVd and HSVd displayed significant differences (Gozmanova et al., 2003). A close look at the region containing RY motifs in PSTVd showed that there is a C-loop (loop 26)(Figure 2A). C-loop is an asymmetric internal loop, which has the following characteristic features: 1) the first base in the longer strand is often a C with some exceptions; 2) the longer strand has two bases forming non-WC base pairs with bases in the other strand; 3) bases from two strands form two triads; 4) this motif often resides in hairpin stem-loop structure (Lescoute et al., 2005; Drsata et al., 2017). Interestingly, our preliminary analysis showed that replacing the C-loop with WC-WC base pairs abolished PSTVd nuclear localization in in situ hybridization analysis.

**Figure 2.**
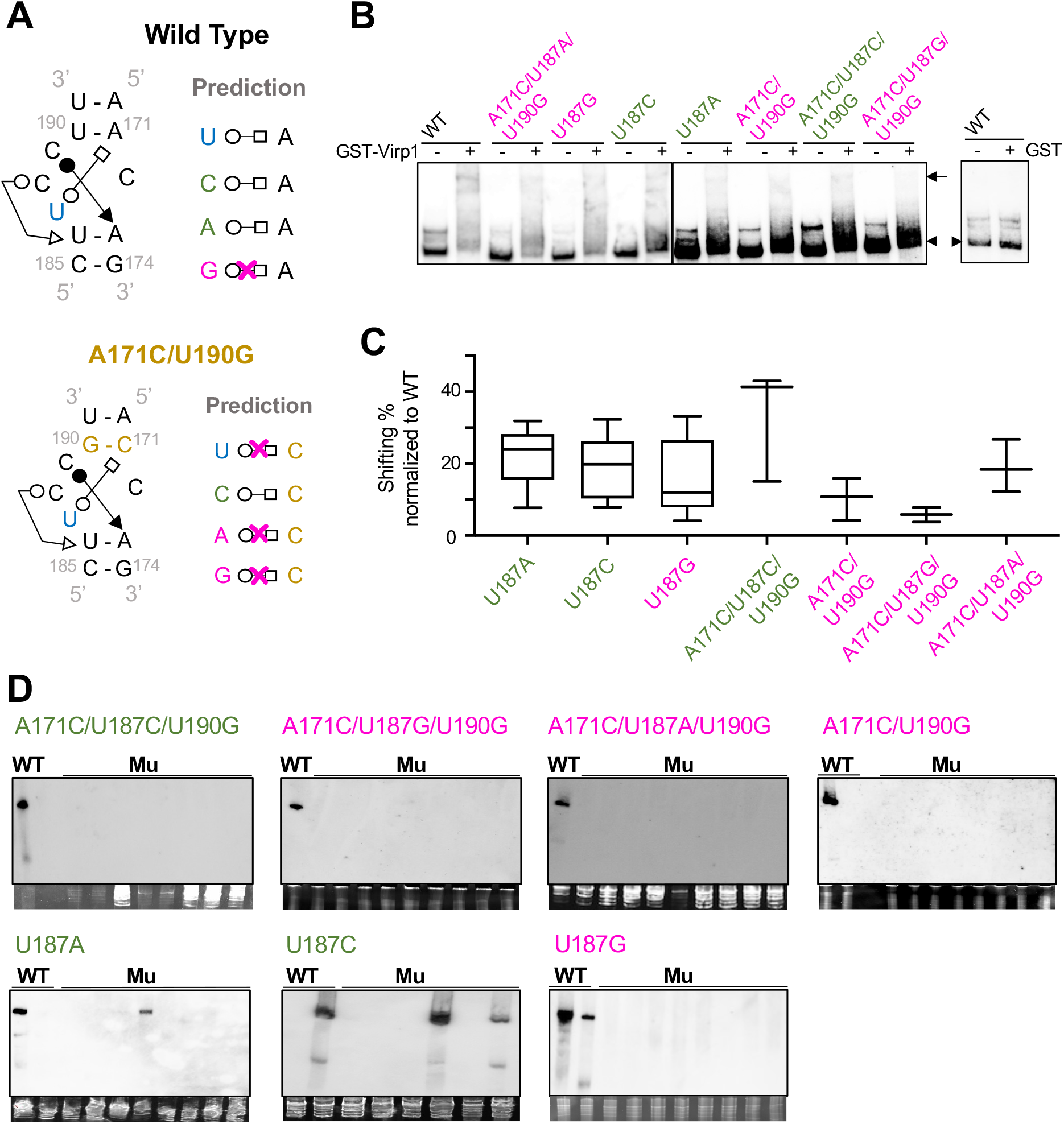
Characterizing PSTVd C-loop using mutagenesis. (A) Rational for C-loop mutant designs. Red crosses indicate that those pairings cannot form in theory. (B) EMSA illustrating the interaction between C-loop mutants and Virp1. (C) Box plot showing quantification of EMSA results that infer the relative affinities of mutant RNAs to Virp1 as compared to that of WT RNA. (D) RNA gel bots detecting the PSTVd systemic infection in *N. benthamiana*. WT PSTVd serves as positive control. Ethidium bromide staining of rRNAs serve as loading control.

According to the C-loop model, PSTVd loop 26 is defined by two WC-WC base pairs (A171-U190 and A173-U186) on both ends. Within this potential C-loop, C189-A173 can form a cis-WC-Sugar base pair (cWS) and U187-A171 can form a trans-WC-Hoogsteen (tWH) base pair. The C189-A173 and U187-A171 base pairs, together with the WC-WC base pairs on both ends, can form two triads (Supplemental Figure 1). C188 and U186 may form a trans-WC-Sugar base pair (tWS) as found in some but not all C-loop structures (Lescoute et al., 2005; Drsata et al., 2017). C172 is predicted as a free-standing base that is not involved in any base-pairing. This PSTVd C-loop model is well supported by the chemical mapping data (Supplemental Figure 1)(Xu et al., 2012; Lopez-Carrasco and Flores, 2017; Steger, 2017). Selective 2′Hydroxyl Acylation analyzed by Primer Extension (SHAPE) assays from multiple studies using different chemicals collectively showed that C172 is highly reactive to modification in vitro and in vivo (Supplemental Figure 1), indicating that it is not involved in base-pairing. In contrast, C189 consistently showed low reactivity in both in vitro and in vivo mapping assays (Supplemental Figure 1), indicating that it is involved in base-pairing. U187 showed medium reactivity in some of the mapping assays but low reactivity in others, which may be attributed to the loop “breathing” effect (Homan et al., 2014). In fact, this is supported by the observation that the partner of U187, A171, also showed relatively high reactivity in some mapping experiments (Supplemental Figure 1). In summary, extensive chemical mapping experiments essentially support that PSTVd loop 26 is a C-loop.

We employed mutational analyses to further test whether loop 26 is a C-loop. Within the PSTVd C-loop (Figure 2A), the cWS base-pairing between C189 and A173 as well as the tWS base-pairing between C188 and U186 are flexible for any nucleotide substitution in theory according to the RNA Basepair Catalog (Stombaugh et al., 2009), so mutations in these two base pairs may not lead to any conclusive result. Instead, we designed substitutions to replace U187 that may or may not maintain similar tWH base-pairing with A171. Alternatively, we replaced the U190-A171 cis-WC-WC base pair with G190-C171. Under this condition, U187 can only be substituted by C187 to maintain the tWH interaction with C171 according to the RNA Basepair Catalog. Using these mutational variants, we performed electrophoretic mobility shift assays (EMSAs) using recombinant Virp1. Interestingly, Virp1 only displayed a strong binding to WT PSTVd in EMSA (Figure 2B and 2C). Among all the tested variants, all structure-maintaining variants have relatively stronger binding to Virp1 as compared with structure-disruptive variants (Figure 2C).

### C-loop is critical for infectivity

As shown in Figure 2D, all PSTVd C-loop disruptive variants and one structure-maintaining mutant (A171C/U187C/U190G) failed to systemically infect *N. benthamiana*. All these infection defective mutants have a weaker binding to Virp1. Two structure-maintaining mutants, U187A and U187C, showed systemic infection. A careful analysis of the RNA progeny in the systemic leaves revealed that none of the progeny maintained the original sequences as inoculum (Supplemental Table 2). Nevertheless, nuclear localization is the prerequisite to initiating replication before mutations occur. Therefore, our data support that the PSTVd structure-maintaining mutants U187A and U187C probably possess the ability to enter the nucleus. Importantly, the data support that PSTVd loop 26 is a C-loop, because only the variants predicted to maintain the C-loop structure have relatively stronger binding with Virp1 and retain the capacity to initiate replication.

### PSTVd C-loop disruptive mutants cannot enter the nucleus

We then analyzed the local leaves inoculated with C-loop variants via Whole-Mount in situ hybridization because it has been well established that viroid-specific riboprobe can highlight infected nuclei thanks to the high concentration of viroid RNAs (Zhu et al., 2002; Qi et al., 2004; Zhong et al., 2007; Takeda et al., 2011). As shown in Figure 3, PSTVd nuclear accumulation was not detected in local leaves inoculated with C-loop structure-disruptive variants (i.e., U187G, A171C/U190G, A171C/U187A/U190G, A171C/U187G/U190G). The structure-maintaining mutants (U187A and U187G) were not included because we cannot distinguish the original inoculum and replication products with mutations in Whole-Mount in situ hybridization assay. In contrast, WT PSTVd formed strong signals in the nucleus (Figure 3). In addition, the replication-defective A261C mutant of PSTVd, which still has nuclear import ability (Zhong et al., 2006), showed detectable nuclear accumulation as well (Figure 3). The nuclear accumulation signal of A261C in Whole-Mount in situ hybridization demonstrated that this assay is sensitive enough to capture imported inoculum without replication. The lack of signal is unlikely caused by RNA stability of C-loop variants, as we often observed C-loop variant inoculums in the local leaves 10 days post-infection. To further test RNA stability, we used agroinfiltration to deliver the cDNAs of C-loop variants into plants. We detected their accumulations comparable to WT transcripts and stronger than the A261C transcripts (Supplemental Figure 2). Altogether, the Whole-Mount in situ hybridization results supported that the C-loop structure-disruptive variants lost their nuclear import ability.

**Figure 3.**
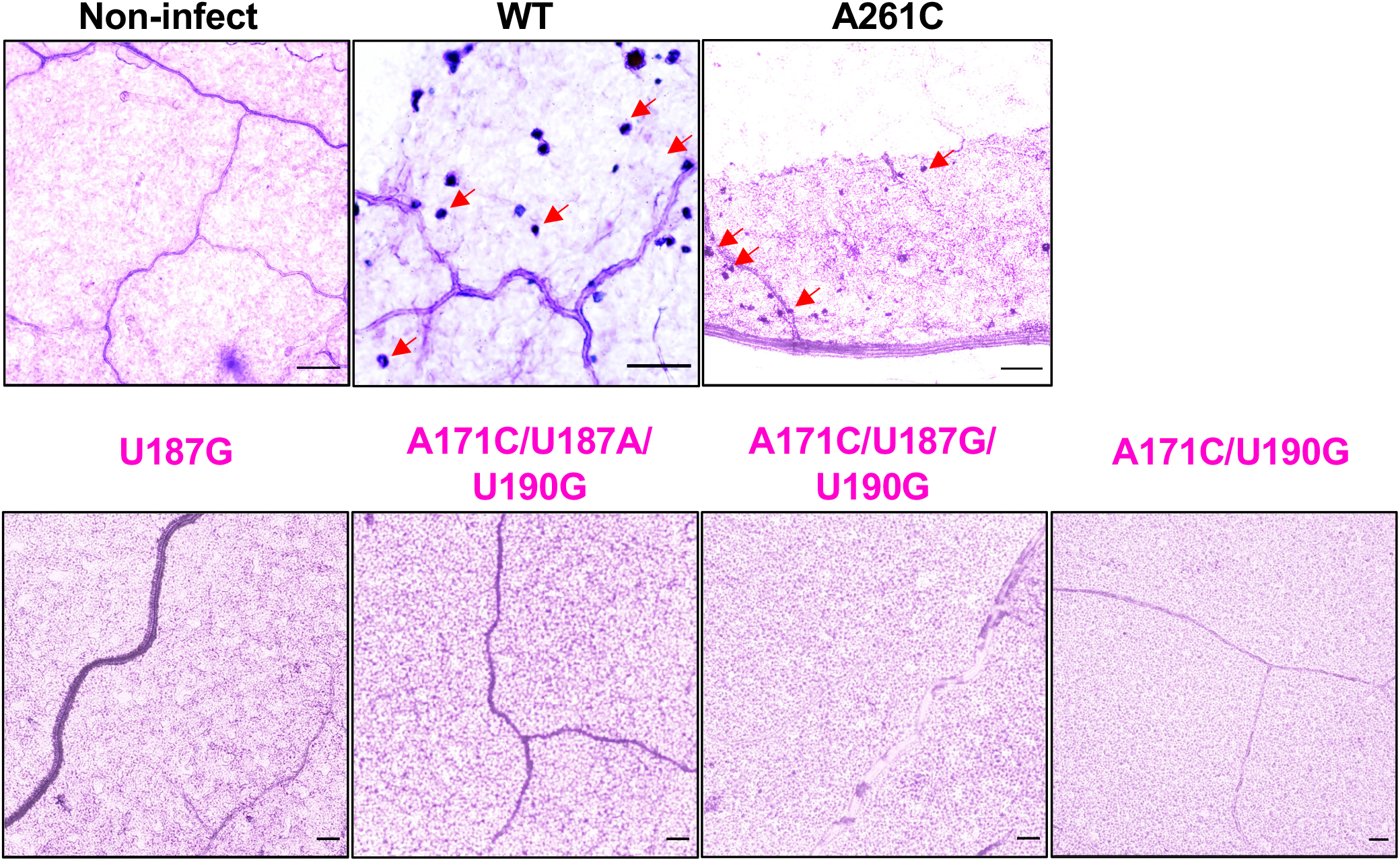
C-loop structure-disruptive mutants failed to accumulate in the nucleus. The purple signals in the whole-mount in situ hybridization show the presence of viroid RNAs in the nuclei. There was no signal in non-infected samples or samples infected with C-loop structure-disruptive mutants, in contrast to the samples infected by WT and A261C mutant. Scale bars, 109 μm. Red arrows indicate a few examples of PSTVd-accumulated nuclei.

### C-loop widely exists in nuclear-replicating viroids

Notably, C-loop can be found in 27 out of 28 formal members and three candidate members of the family *Pospiviroidae* (Supplemental Figure 3). Based on sequence variations and genomic coordination, we can categorize those viroids into two groups (Supplemental Figure 3). Interestingly, there are 11 viroids, including PSTVd, containing exactly the same C-loop with identical genomic localization. The remaining 19 viroids in Supplemental Figure 3 have C-loop structures with diverse sequence variations and genomic localization patterns, which still fit the C-loop model. This observation demonstrates that C-loop is likely a common motif exploited by viroids for nuclear import.

Notably, we also found a variant version of C-loop in HSVd (Figure 4A), a PSTVd relative that has a slightly weaker binding to Virp1 (Maniataki et al., 2003). To test this C-loop variant, we replaced the C128-G171 cis-WC-WC base pair with G-C, A-U, or U-A (Figure 4A). Only the A128-U171 substitution is predicted to disrupt the tWH base pair within the C-loop. Again, all HSVd C-loop variants, including one structure-disruptive and two structure-maintaining variants, exhibited much-reduced binding to Virp1 (Figure 4B and 4C). Relatively, both structure-maintaining variants exhibited a slightly stronger binding to Virp1 as compared with the structure-disruptive variant. Since we observed reduced binding in all the mutational designs, one more mutant (G157C), which affects an adjacent loop to the C-loop in HSVd, was inclcuded as a control. This mutant now binds to Virp1 comparably to the binding between ViRP1 and WT HSVd (Figure 4B and 4C).

**Figure 4.**
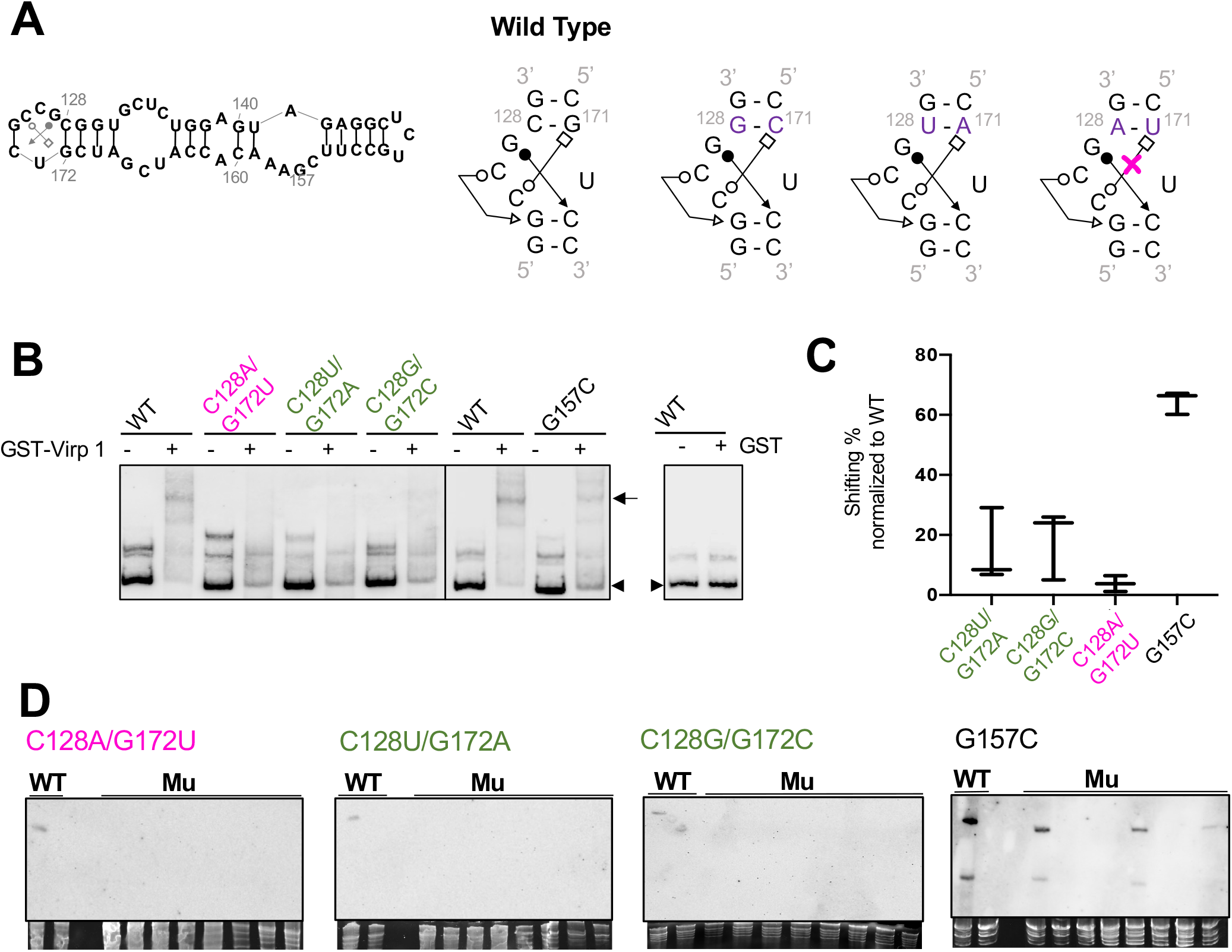
Characterizing a C-loop variant in HSVd. (A) Rational for C-loop mutant designs. (B) EMSA illustrating the interaction between C-loop mutants and Virp1. (C) Box plot showing quantification of EMSA results that infer the relative affinities of mutant RNAs to Virp1 as compared to that of WT RNA. (D) RNA gel bots detecting the HSVd systemic infection in *N. benthamiana*. WT HSVd serves as positive controls. Ethidium bromide staining of rRNAs serves as loading control.

When we used HSVd C-loop variants and the G157C mutant to infect *N. benthamiana* plants, only G157C can accomplish successful infection (Figure 4D). Sequencing of the progeny confirmed that the G157C mutation was retained in the progeny in systemic leaves (Supplemental Table 2). Altogether, our observation supports that C-loop is critical for HSVd infectivity and Virp1 specifically recognizes HSVd C-loop. Since our structure-maintaining mutants showed weak binding to Virp1, it may prefer certain nucleotides in the composition of the C-loop.

## Discussion

Proper subcellular localization dictates the function of biomolecules, including various cellular and infectious RNAs. While a majority of cellular RNAs are made in the nucleus and then either stay in the nucleus or are transported to the cytoplasm for function, more and more RNAs were found to traffic in a reverse direction from the cytoplasm to the nucleus participating in diverse biological processes (Rudt and Pieler, 1996; Chou et al., 1998; Gao et al., 2012; Ye et al., 2012; Kramer and Hopper, 2013; Chaturvedi et al., 2014; Long et al., 2021). However, the mechanism underlying RNA nuclear import is poorly understood. Here, we present evidence unraveling an IMPa-4-based cellular pathway, together with cellular protein Virp1, in transporting pathogenic noncoding RNAs (i.e., viroids) from the cytoplasm to the nucleus in plants (Figure 5). Notably, we identified one genetic element, an RNA C-loop, as a critical signal for viroid RNA nuclear import. PSTVd C-loop model is supported by chemical mapping data (Supplemental Figure 1) and our mutational analyses. Disrupting C-loop resulted in much-reduced Virp1-binding ability, loss of infectivity, and the lack of nuclear accumulation signal in inoculated leaves. The absence of nuclear signal in Whole-Mount in situ hybridization using C-loop mutant inoculated samples is unlikely attributable to RNA stability because C-loop mutant RNAs have comparable stability in plants as wild type RNAs (Supplemental Figure 2).

**Figure 5.**
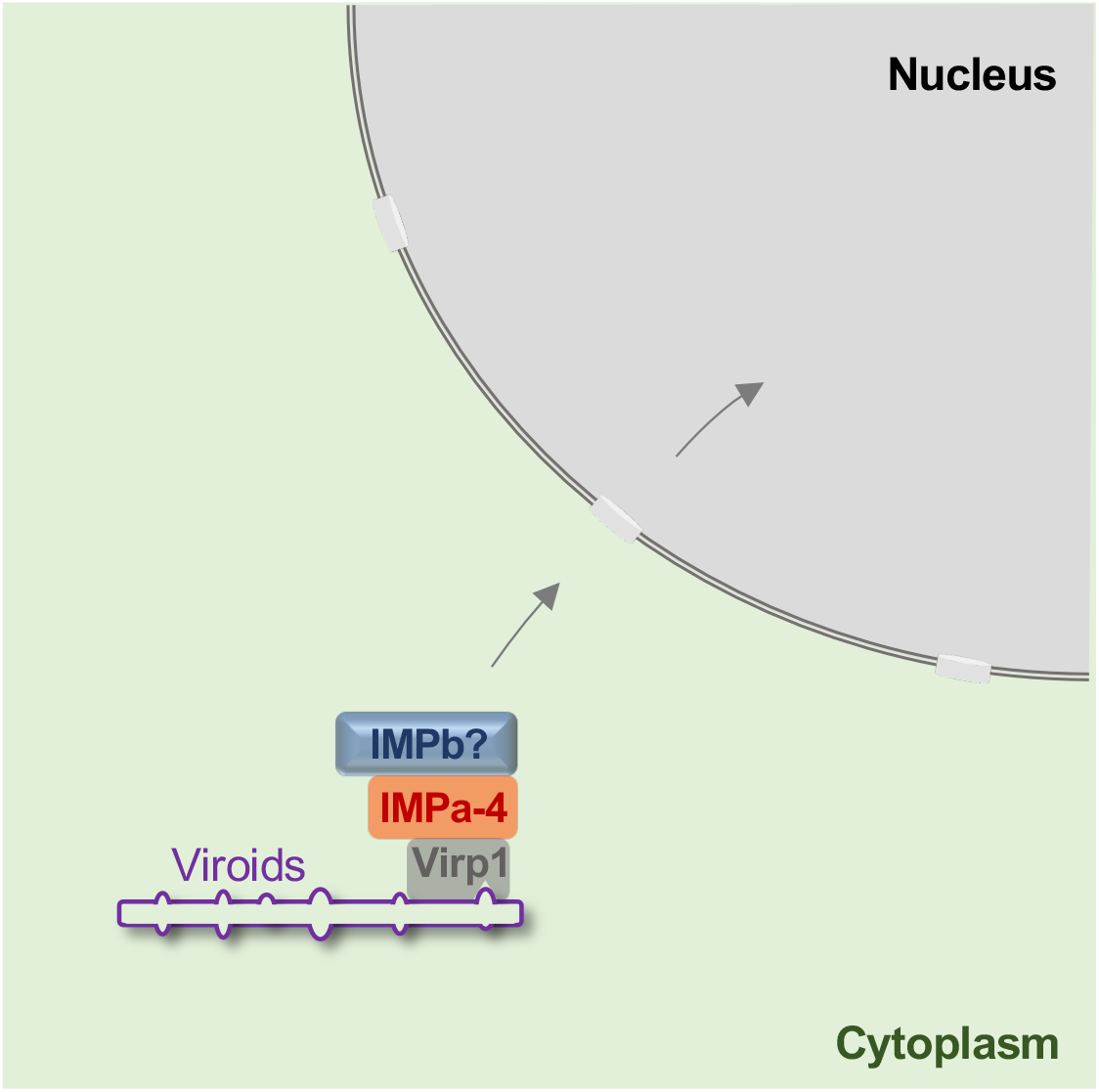
A working model illustrating the IMPa-4/Virp1/C-loop based RNA nuclear import.

Previous studies suggest that Virp1 recognizes RY motifs in viroids (R: A or G; Y: C or U) (Gozmanova et al., 2003; Maniataki et al., 2003). RY motif and C-loop partially overlap in some viroids, like PSTVd. The drastic changes in binding and infectivity of point mutations that disrupt PSTVd C-loop support the strong role of C-loop for Virp1 recognition. More importantly, HSVd C-loop disruptive mutants that are not overlapping with RY motifs have a strong effect on infectivity and Virp1-binding. In contrast, the G157C mutant that overlaps with the RY-motif retains infectivity and Virp1-binding ability. Altogether, our data strongly support C-loop as a bona fide signal for selective nuclear import of RNA.

C-loop has been found in many rRNAs (Lescoute et al., 2005; Drsata et al., 2017), a bacterial mRNA (Torres-Larios et al., 2002), and some conserved mammalian noncoding RNAs (Iacoangeli and Tiedge, 2013). In general, C-loop increases the local helical twist of RNA helices (Afonin and Leontis, 2006). C-loop in the mRNA and those mammalian noncoding RNAs are involved in translational regulation (Torres-Larios et al., 2002; Iacoangeli and Tiedge, 2013). Our data expand the function of the C-loop and uncover Virp1 as a new protein partner for this RNA motif.

Virp1 was first identified as a PSTVd interacting protein through screening cDNA library from PSTVd-infected tomato for RNA ligand binding (Martinez de Alba et al., 2003). Early work suggests that Virp1 is important for PSTVd systemic trafficking because the binding-deficient PSTVd mutants failed to achieve systemic trafficking (Maniataki et al., 2003). Later on, protoplast assay using Virp1 down-regulated cells suggested that Virp1 has a role in PSTVd replication (Kalantidis et al., 2007). Very recently, Virp1 has been implied to regulate CEVd nuclear import in an in situ assay based on onion cells (Wu et al., 2019). Our data provide compelling evidence supporting a critical role of Virp1 in viroid nuclear import. Notably, both IMPa-4 and Virp1 have been implicated in T-DNA nuclear import (Crane and Gelvin, 2007; Bhattacharjee et al., 2008). It will be interesting to further investigate whether IMPa-4 and Virp1 also participate in the nuclear import of viral DNAs and/or cellular RNAs in the future.

Currently, there are 28 formal members of the family *Pospiviroidae* (Di Serio et al., 2021), and 27 of them (except citrus dwarfing viroid) possess a C-loop. Eleven viroids, including eight out of nine members in the genus *Pospiviroid* where PSTVd belongs, possess an identical C-loop in their genomes. A close look at the rest of the viroids of the family *Pospiviroidae* suggests the existence of C-loop, with some variations in sequences and genome localization, in all but citrus dwarfing viroid (Supplemental Figure 3). Notably, these viroid genomic structures are supported by SHAPE analyses (Xu et al., 2012; Giguere et al., 2014; Giguere and Perreault, 2017), except for citrus bark cracking viroid and citrus viroid-VI whose structures were predicted using mFOLD (Zuker, 2003). Therefore, a conserved nuclear import signal likely exists in nearly all nuclear-replicating viroids. Interestingly, C-loop variants can also be found in mexican papita viroid, citrus viroid-IV, and grapevine latent viroid, which are candidate members of *Pospiviroidae*. Future functional investigation on those C-loop variants can provide insights into the precise structural basis and critical nucleotide preferences in mediating RNA nuclear import. It is also interesting to analyze citrus dwarfing viroid to test 1) whether it possesses an alternative binding site for Virp1 and/or 2) whether there is an alternative nuclear import route.

Interestingly, Q-satRNA appears to have a C-loop in its RNA sequence as well. EMSA testing using a C-Loop disruptive Q-satRNA showed significantly reduced binding to Virp1 as compared with that of WT Q-satRNA, further supporting the critical role of C-loop in binding with Virp1 (Supplemental Figure 4). Therefore, C-loop-based RNA nuclear import is possibly exploited by infectious RNAs in common. Whether any cellular RNA follows this pathway for nuclear import to exert functions in plants deserves future investigation. Our study paves the way to explore RNA nuclear import machinery and outlines a model for structural motif-based RNA subcellular localization. This line of research may lead to a comprehensive understanding of the accurate localization of RNAs in cells and future manipulation of subcellular localizations of various RNAs for functional studies and applications.

## Materials and Methods

### Plant growth

*Arabidopsis* plants were grown in a growth chamber at 22°C with a 10/14 h light/dark cycle. *N. benthamiana* plants were grown in a growth chamber at 25°C with a 14/10 h light/dark cycle. *N. benthamiana* seedlings at the four-leaf stage were inoculated with water or water containing 150 ng of in vitro transcribed viroid RNAs. The viroid infection was analyzed by RNA gel blots using local leaves 10 days post-inoculation and systemic leaves three weeks post-inoculation as described previously. Agroinfiltration followed an established protocol (Wang et al., 2016).

### DNA clones

cDNAs of some Arabidopsis Importin alpha subunits in pC-TAPa or Lic6 vectors were purchased from ABRC (Ohio State University, Columbus, OH): IMPa1 (DKLAT3G06720), IMPa2 (DKLAT4G16143), IMPa3 (DKLAT4G02150), IMPa-4 (DKLAT1G09270), IMPa5 (DKLAT5G49310.1) and IMPa6 (DKLAT1G02690). IMPa7 cDNA in pDONR221 vector (DQ446636) was purchased from ABRC and recombinated into pC-TAPa vector (ABRC) via LR clonase (Thermo Fisher Scientific, Waltham, MA). IMPa8 and IMPa9 were amplified using gene-specific primers (Supplemental Table 3) and cloned into pCR8 (Thermo Fisher Scientific), which were then recombinated into pC-TAPa via LR clonase.

To generate the pTRV2-IMPa-4sil clone, two specific primers (Supplemental Table 3) for *N. benthamiana* IMPa-4 fragment were used for genomic PCR and followed by digestion with *BamH*I and *Xho*I (New England Biolabs, Ipswich, MA). The pTRV2 vector (ABRC) linearized by *BamH*I and *Xho*I were used for ligation with the digested NbIMPa-4 fragments. Since we cannot reach 100% PSTVd infection in *N. benthamiana*, we then decided to use tomato for the VIGS assay. Based on the high sequence homology of IMPa-4 in tomato and *N. benthamiana*, we used the same pTRV2-IMPa-4sil clone for infiltration. Based on the BLAST search using Sol genomics database (https://solgenomics.net), our cloned fragment specifically targets IMPa-4 homologs in tomato and *N. benthamiana*. pTRV2 variants in agrobacterium GV3101 were mixed with Agrobacterium harboring pTRV1 (ABRC) for VIGS infiltration into one cotyledon of tomato seedlings, while the other cotyledon was used for inoculation with PSTVd RNA transcripts. Plants were subjected to RNA gel blot to analyze PSTVd and TRV titers, as well as the expression levels of IMPa-4 and Histone H2A (see Supplemental Table 3 for primer details). The TRV probe was described previously (Zheng et al., 2017a).

Virp1 from Arabidopsis was cloned via RT-PCT using gene-specific primers (Supplemental Table 3). The cloned Virp1 cDNA was inserted in pCR8 vector and then recombinated into pMDC7 vector (modified to include a N-FLAG tag; inherited from Biao Ding at Ohio State University) for agroinfiltration or pDEST15 vector (Thermo Fisher Scientific) for bacterial expression, via LR clonase. GST construct for bacterial expression was a gift from Svetlana Folimonova at University of Florida.

The cDNAs of WT and mutant Q-satRNAs were commercially synthesized (Genscript, Piscataway, NJ). The cDNAs were amplified (see Supplemental Table 3 for primer sequences) and ligated into pGEM-T vector (Promega, Madison, WI). To generate RNA substrates for EMSA, *Spe*I (New England Biolabs) linearized plasmids (pGEMT-Q-satRNA^WT^ and pGEMT-Q-satRNA^mu^) were subject to *in vitro* transcription using T7 MEGAscript kit (Thermo Fisher Scientific).

To generate RNA inocula, pRZ:Int construct (Wang et al., 2007) was by *Hind*III (New England Biolabs) followed by *in vitro* transcription using T7 MEGAscript kit. pT3:HSVd^RZ^ (Tu HSVd2-7 in the 83-82 orientation) used pGEM-T vector with the insertion cloned from HSVd-RZ (a gift from Dr. Robert Owens at USDA-ARS) via T3-HSVd-f and RZ-r primers (Supplemental Table 3). pT3:HSVd^RZ^ was linearized by *Hind*III followed by *in vitro* transcription using T3 MEGAscript kit (Thermo Fisher Scientific). All RNA *in vitro* transcripts were purified using the MEGAclear kit (Thermo Fisher Scientific).

To generate Riboprobes, pInt(-) (Zheng et al., 2017b) was linearized by *Spe*I (New England Biolabs) as the template and T7 MAXIscript kit (Thermo Fisher Scientific) was used to generate probe. pHSVd-monomer was based on pGEM-T vector (Promega) via insertion of HSVd cDNA cloned from HSVd-RZ plasmid via HSVd-f and HSVd-r primers (Supplemental Table 3). The pHSVd-Monomer was linearized by *Nco*I (New England Biolabs) as the template and SP6 MAXIscript kit (Thermo Fisher Scientific) was used to generate probe. To generate Q-satRNA probe, pGEMT-Q-satRNA^WT^ was linearized by *Nco*I (New England Biolabs) and subject to in vitro transcription using SP6 MAXIscript kit.

To generate WT, A261C, and C-loop mutant constructs for agroinfiltration, the corresponding pRZ:Int plasmids harboring the correct PSTVd cDNAs served as templates for PCR (using RZ-f and RZ-r primers; see Supplemental Table 3 for primer sequences). The PCR products were inserted into pENTR-D-TOPO vector (Thermo Fisher Scientific). The series of pENTR-RZ:Int plasmids were recombinated into CD3-1656 (ABRC) via LR clonase. The CD3-1656-RZ:Int plasmid series were transformed into Agrobacterium strain GV3101 for agroinfiltration.

All the constructs have been verified using Sanger sequencing.

### RNA immunoprecipitation

RNA-immunoprecipitation (RIP) was performed according to a previously described protocol (Wang et al., 2016) with minor modifications. Briefly, PSTVd-infected leaves were harvest 3 days post agroinfiltration of Importin alpha cDNAs. The cell lysates were incubated with magnetic mouse IgG beads (Cell Signaling, Danvers, MA) for 2 h at 4°C. The input lysate and purified fractions were subject to immunoblotting and RT-PCR (after RNA purification). The primers for detecting PSTVd and Histone 2A mRNA were listed in Supplemental Table 3.

### Co-Immunoprecipitation

Co-immunoprecipitation was following a recent report (Chen et al., 2020) with minor modifications. FLAG-tagged Virp1 was co-expressed with MYC-tagged IMPa1 or IMPa-4 via agroinfiltration. Three days post infiltration, 4 mM 17-b-estradiol was infiltrated in leaves 6 h before sampling. The cell lysates from leaf samples were incubated with anti-FLAG antibody for 1 h at 4°C. The magnetic protein A/G beads (Thermo Fisher Scientific) were then added to the lysate for another 1 h incubation at 4°C with mild shaking. The beads were washed twice with 1X PBST buffer (137 mM NaCl, 2.7 mM KCl, 10 mM Na_2_HPO_4_, 1.8 mM KH_2_PO_4,_ 0.1% Triton X-100) and once with distilled water. The bound proteins were eluted using IgG elution buffer (Thermo Fisher Scientific) and then subject to immunoblots.

### Protein purification

GST and Recombinant Virp1-GST proteins were expressed in *Escherichia coli* Rosetta strain (EMD Millipore, Burlington, MA). Cells were grown overnight at 37°C in LB media supplied with ampicillin (0.1 mg/ml) and chloramphenicol (0.034 mg/ml). An aliquot of cells with OD600 = 0.1 was inoculated into fresh LB supplied with antibiotics the next day. Once the cell density (OD600) reached 0.5-0.7, 0.4 mM IPTG (final concentration) was added to the culture to induce protein expression. After inducing at 20°C overnight, 100 ml culture was harvested by centrifugation at 8,000 rpm for 8 min. Pellets were re-suspended in 1X PBS buffer (137 mM NaCl, 2.7 mM KCl, 10 mM Na_2_HPO_4_, and 1.8 mM KH_2_PO_4_) supplement with 20 mM PMSF and sonicated to lyse the cells. The cell lysate was then centrifuged at 10,800 rpm for 30 min at 4°C. The supernatant was collected and incubated for 1 h with 2 ml of 50% slurry of Glutathione Resin (Genscript) before loading onto an empty EconoPac gravity-flow column (Bio-Rad Laboratories, Hercules, CA). The resin was then washed with 10 mL 1xPBS followed by applying 10 mL elution buffer (50 mM Tris-HCl pH 8.0 and 10 mM reduced glutathione). The elutes were concentrated using an Amicon protein concentrator (MilliporeSigma, Burlington, MA). Proteins were then separated by 8% SDS-PAGE electrophoresis followed by Coomassie blue staining and de-staining to estimate concentration using a BSA standard as reference.

### Electrophoresis mobility shifting assays (EMSAs)

The detailed protocol has been reported previously (Gozmanova et al., 2003). Binding assays that contained RNA in the absence or presence of different amounts of GST or Virp1-GST proteins were incubated at 28°C for 30 min. The binding buffer was composed of 10 mM HEPES-NaOH (pH 8.0), 50 mM KCl, 100 mM EDTA, and 5% glycerol. Electrophoresis for the binding assay was performed on ice in 6% polyacrylamide (29:1) gels at 140 V using 0.5X TBE (50 mM Tris, 50 mM boric acid, 1 mM EDTA, pH 8.3) for 1.6 h. The following steps are described below in the RNA gel blots section. The percentage of shifted variant RNAs was normalized to that of WT RNAs to infer a relative binding strength to Virp1, based on at least three replicates.

### Tissue processing and in situ hybridization

The tissue fixation and processing were largely described previously (Takeda et al., 2011) with minor modification. Briefly, 8 DPI leaf samples were fixed in FAA solution (50% ethanol/5% formaldehyde/5% acetic acid) for 30 min and then dehydrated by a step-wise gradient of ethanol solutions. The samples were washed by 1XPBS and treated with 10 mg/mL of proteinase K for 20 min at 37°C. Then, the samples were hybridized with Dig-labeled antisense riboprobes (generated as above-mentioned) at 50°C overnight. The samples were washed, incubated with anti-DIG monoclonal antibody (MilliporeSigma) and NBT/BCIP substrate (MilliporeSigma) subsequentially, and mounted with Permount (Thermo Fisher Scientific) for visualization using an Olympus CX23 light microscope. The scale bars were calculated using ImageJ (https://imagej.nih.gov/ij/).

### RNA gel blots and immunoblots

After electrophoresis, RNAs were then transferred to Hybond-XL nylon membranes (Amersham Biosciences, Little Chalfont, United Kingdom) via a semi-dry transfer cassette (Bio-Rad Laboratories) and were immobilized by a UV-crosslinker (UVP, Upland, CA). RNAs were then detected by DIG-labeled UTP probes. AP-conjugated anti-DIG monoclonal antibody was used in combination with Immun-Star substrates (Bio-Rad Laboratories). Signals were captured by ChemiDoc (Bio-Rad Laboratories).

After SDS-PAGE electrophoresis, we followed the previously described protocol for immunoblotting (Jiang et al., 2019). IMPas were detected by a monoclonal mouse anti-Myc antibody (MilliporeSigma; 1:3,000 dilution). Virp1 was detected by a monoclonal mouse anti-FLAG antibody (Thermo Fisher Scientific; 1:1,000 dilution). HRP-conjugated anti-mouse serum (Bio-Rad Laboratories) was diluted at 1:2000. SuperSignal West Dura (Thermo Fisher Scientific) was used as the substrate. Signals were captured by ChemiDoc (Bio-Rad Laboratories).

## Data availability

The published RNA-Seq dataset has been deposited in the NCBI SRA with accession number SRP093503.

## Acknowledgements

This work was supported by grants from the US National Science Foundation (MCB-1906060 to YW) and the US National Institute of General Medical Sciences (1R15GM135893 to YW). We are indebted to Robert Owens (USDA-ARS) and Svetlana Folimonova (University of Florida) for their gifts of DNA constructs. We are grateful for the constructive comments from Donna Gordon (Mississippi State University). We thank Laxmi Kharel (Mississippi State University) for technical support. This work is dedicated to the late Professors Biao Ding (Ohio State University) and Neocles B. Leontis (Bowling Green State University), who participated at the inception stage.

## Author contributions

RT and YW conceived the study. YW designed the experiments. JM, SDM, and YW performed the experiments. JM and YW analyzed the data. YW wrote the manuscript. MJ, WJP, MW, BL, and YW discussed the results and revised the manuscript.

