## Supplemental Figure 1 for "A nuclear import pathway exploited by pathogenic noncoding RNAs"

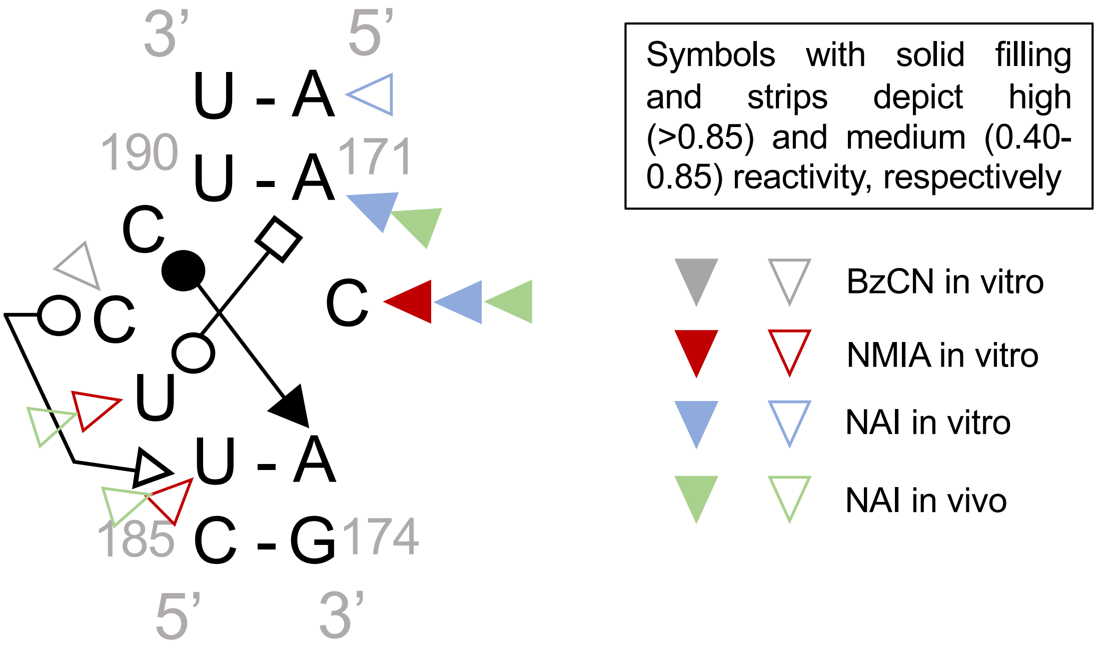


**Supplemental Figure 1** Selective 2’ Hydroxyl Acylation analyzed by Primer Extension (SHAPE) analyses support PSTVd C-loop model. The figure plotted using the published data (Xu et al., 2012; Lopez-Carrasco and Flores, 2017). Bases with low reactivities were not highlighted. BzCN, Benzoyl Cyanide. NMIA, N-methylisatoic anhydride. NAI, 2-methylnicotinic acid imidazolide.
