## Supplemental Figure 2 for "A nuclear import pathway exploited by pathogenic noncoding RNAs"

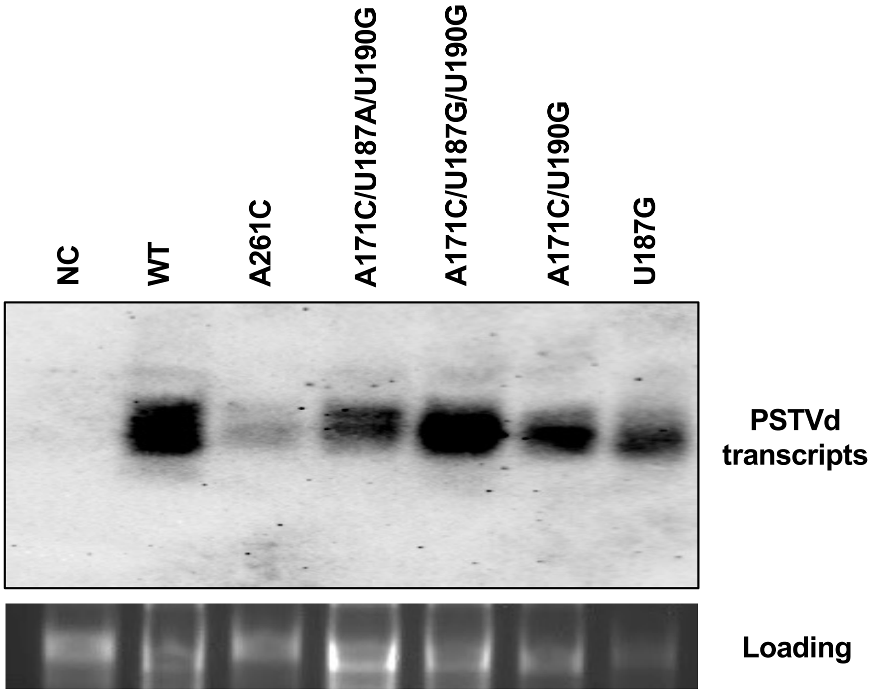


**Supplemental Figure 2** The RNA stability of PSTVd C-loop variants. C-loop mutants, in comparison to wildtype (WT) and A261C mutant, were expressed in *N. benthamiana* via agroinfiltration using CaMV 35S promoter driven RZ:Int-based constructs. Total RNAs from 4 days post infiltrated leaves were run in 2% agarose gel and blotted with PSTVd specific ribo-probes. Ethidium bromide staining of rRNAs serves as loading control. NC, negative control.
