## Supplemental Figure 3 for "A nuclear import pathway exploited by pathogenic noncoding RNAs"

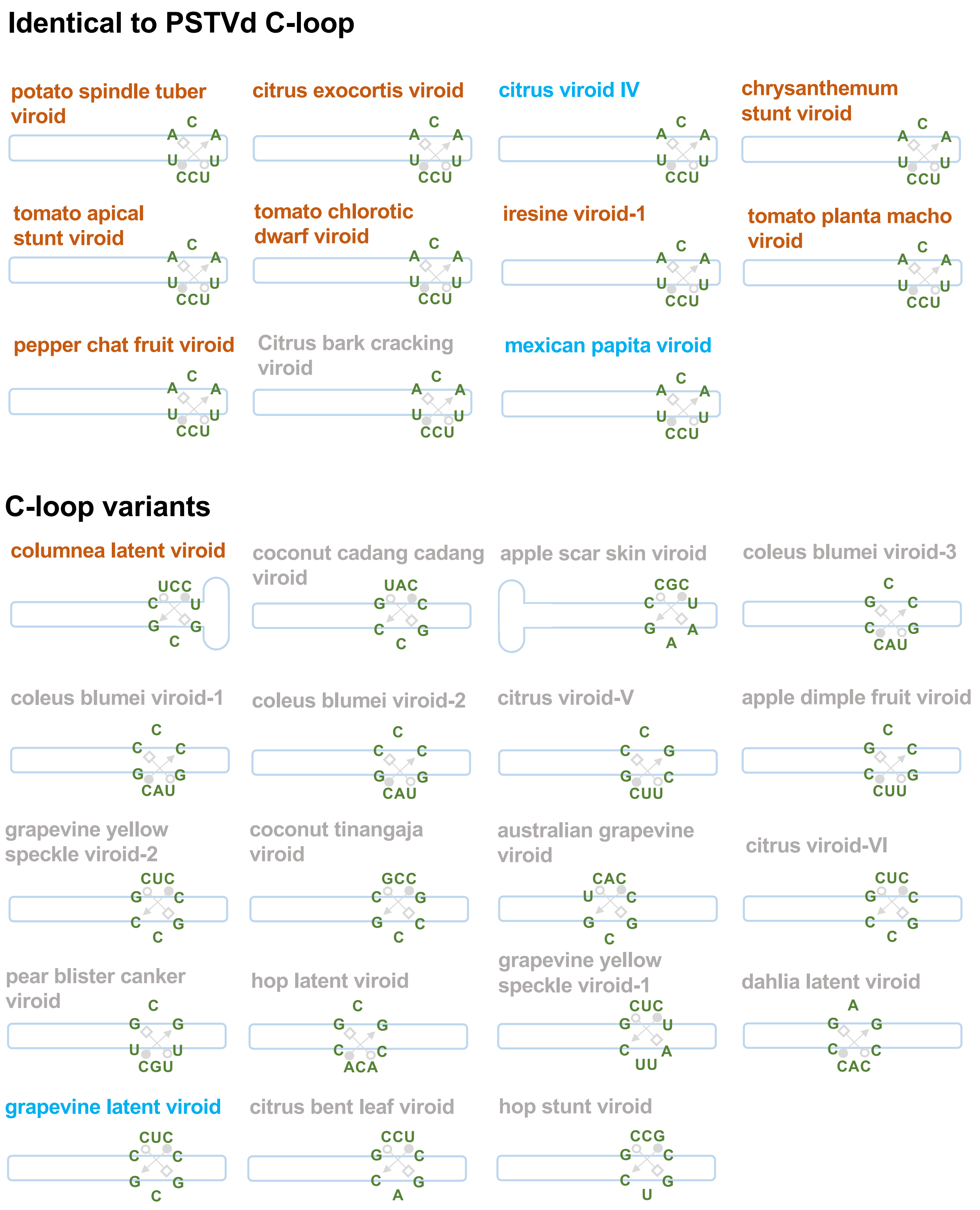


**Supplemental Figure 3** C-loop in nuclear-replicating viroids. Illustration of viroid C-loop sequences and relative genomic loci. Viroids whose names are in brown belong to the genus *Pospiviroid*. Viroids whose names in blue were considered but not confirmed as members of the family *Pospiviroidae* in the latest taxonomy.
