## Supplemental Figure 4 for "A nuclear import pathway exploited by pathogenic noncoding RNAs"

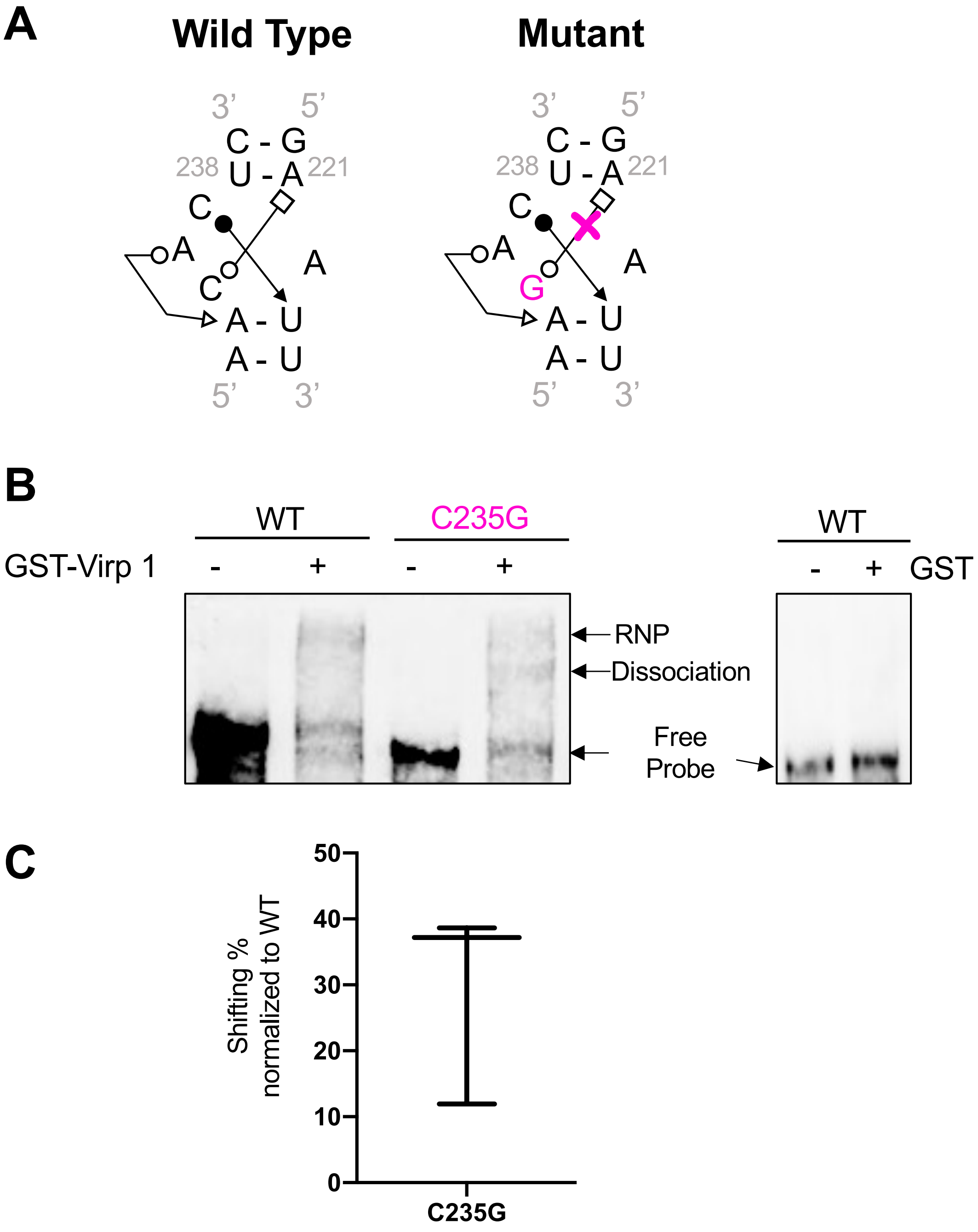


**Supplemental Figure 4** ViRP1 interaction with Q-satRNA. (A) Rational for C-loop mutant designs. (B) EMSA illustrating that C-loop disruptive mutant C235G significantly reduced Q-satRNA binding with Virp1. (C) Box plot showing quantification of EMSA results that infer the relative affinity of C235G RNA to Virp1 as compared to that of WT RNA.
