## Supplemental Table 1 for "A nuclear import pathway exploited by pathogenic noncoding RNAs"

**Supplemental Table 1.** IMPa homologs in tomato. The normalized reads (FPKM) of tomato IMPas in RNA-Seq dataset are listed. NA, not found.

| | | | Mock vs PSTVd | | | | | | | | | |
|---|---|---|---|---|---|---|---|---|---|---|---|---|
| | Arabidopsis | Tomato | **Mock_rep1** | **Mock_rep2** | **Mock_rep3** | **mean** | **PSTVd_rep1** | **PSTVd_rep2** | **PSTVd_rep3** | **mean** | **ratio** | **adjust p** |
| IMPa1 | AT3G06720 | Solyc08g041890.4.1 | 37.26 | 28.56 | 33.3 | 33.04 | 43.64 | 42.3 | 33.43 | 39.79 | 1.2 | 1 |
| IMPa2 | AT4G16143 | Solyc01g060470.3.1 | 37.78 | 31.92 | 33.94 | 34.55 | 49.3 | 69.71 | 59.71 | 59.57 | 1.72 | 0.065823 |
| IMPa3 | AT4G02150 | Solyc06g009750.4.1 | 21.24 | 10.49 | 25.31 | 19.01 | 19.84 | 30.67 | 27.4 | 25.97 | 1.37 | 0.532009 |
| IMPa4 | AT1G09270 | Solyc01g100720.3.1 | 55.79 | 43.04 | 52.85 | 50.56 | 51.12 | 75.54 | 79.34 | 68.67 | 1.36 | 0.230871 |
| IMPa5 | AT5G49310 | NA | | | | | | | | | | |
| IMPa6 | AT1G02690 | NA | | | | | | | | | | |
| IMPa7 | AT3G05720 | NA | | | | | | | | | | |
| IMPa8 | AT5G52000 | NA | | | | | | | | | | |
| IMPa9 | AT5G03070 | Solyc10g084270.2.1 | 27.93 | 26.51 | 27.78 | 27.41 | 29.09 | 24.88 | 12.24 | 22.07 | 0.81 | 0.337192 |
