## Supplemental Table 2 for "A nuclear import pathway exploited by pathogenic noncoding RNAs"

**Supplemental Table 2.** PSTVd and HSVd progeny in systemic leaves.

| Inoculum | Progeny in systemic leaves | Count |
|---|---|---|
| PSTVd U187A | WT | 6 |
| | U252A | 1 |
| | U24C | 1 |
| | U240A | 1 |
| | U27C/C102U/C117U/A274G/U356C | 1 |
| PSTVd U187C | WT | 6 |
| | U187C/A119G | 1 |
| | G287A | 1 |
| | C117U | 1 |
| | A152G | 1 |
| HSVd G157C | G157C | 5 |
| | G157C/U258C | 1 |
| | G157C/A158G/G239A | 1 |
