## Supplemental Table 3 for "A nuclear import pathway exploited by pathogenic noncoding RNAs"

**Supplemental Table 3.** Primer sequences. Nb, Nicotiana benthamiana. Sl, Solanum lycopersicum (tomato).

| | Primer name | Sequences | |
|---|---|---|---|
| Q-satRNA | f | aggtctcacatgttttgtttgttagagaattgcgtagagggg | |
| | r | tggtctctggccgggtcctggtagggaatgataaac | |
| HSVd | T3-HSVd-f | ggggacaagtttgtacaaaaaagcagaattaaccctcactaaaggcaactcttctcaga | |
| | RZ-r | cgggtaccaggtaatataccacaac | |
| | HSVd-f | gcaactcttctcagaatccagcg | |
| | HSVd-r | cccggggctcctttctcag | |
| PSTVd | 95f | ggggaaacctggagcgaactgg | |
| | 94r | cccggggatccctgaagcgctcc | |
| Histone H2A | Nb f | atggatactagcggcaaagcgaag | |
| | Nb r | ctaagccttcttaggagatttggtag | |
| | RTr | cgagaacagcagccaagtaaacg | |
| | Sl f | atggagtctaccggaaaagtgaag | |
| | Sl r | tgccttcttgggagatttggtag | |
| IMPa8 | f | atggcttggaaaacagaggtgaacga | |
| | r | cacctgaaagtccacatcatcacatc | |
| IMPa9 | f | atggcggatgatggctccgcct | |
| | r | ttcatcgattccataatcttcaccaaagtatttatc | |
| ViRP1 | f | atggctccggctgttttcgctac | |
| | r | ttaacattgggcttcttttgcttccac | |
| NbIMPa-4 | BamHI p f | aaggatcccttcgacccggcactcg | Note: for generating TRV2-IMPa4SiI |
| | XhoI p r | aaactcgagcctttttctcaatagtggcaggt | |
| SlIMPa-4 | p f | gctacctctggaggatctaatga | |
| | p r | gaacattaggctggttgtttccg | |
| RZ:Int | RZ-f | caccggaattcgagctcggtacccgg | |
| | RZ-r | cgggtaccaggtaatataccacaac | |
